# Differences in isotopic compositions of individual grains and aggregated seed samples affect interpretation of ancient plant cultivation practices

**DOI:** 10.1101/2024.08.15.607704

**Authors:** Nathaniel F. James, Christine Winter-Schuh, J. Mark Kenoyer, Jade D’Alpoim Guedes, Cheryl A. Makarewicz

## Abstract

The stable carbon (δ^13^C) and nitrogen (δ^15^N) isotope analysis of charred archaeological grains provides a remarkably precise scale of information: the growing conditions under which a plant was cultivated in a single field and season. Here we investigate how the measurement of single individual grains or aggregate ‘bulk’ samples for carbon and nitrogen isotopes impacts how we characterize variation and, consequently, our interpretations of ancient cultivation practices. Using experimentally grown barley (*Hordeum vulgare* var. *nudum)*, this work investigates δ^13^C and δ^15^N intra-panicle variation between both uncharred and charred individual grains from four plants. We found limited intra- and inter-panicle isotopic variation in single grain isotope values, ca. 0.5‰ in δ^13^C and ca. 1‰ in δ^15^N, reemphasizing the degree to which grains are representative of their local growing conditions. To explore the interpretive impact of aggregate versus single-grain isotopic sampling, we measured charred barley recovered from a single storage context excavated from Trench 42 (ca. 1900 BCE) at Harappa. Aggregate samples of a random selection of Trench 42 barley demonstrated remarkable inter-sample homogeneity, with a less than 0.5‰ difference in δ^13^C and δ^15^N values, reinforcing the ability of aggregate samples to capture a representative isotopic average of a single depositional context. However, the measurement of single-grains revealed moderate 2–3‰ variation in δ^13^C, and an outstandingly wide isotopic variation of ca. 8‰ in δ^15^N values, indicating the degree to which cultivation practices varied beyond what the bulk samples indicated. These results highlight how decisions in the selection and measurement of archaeological grains for isotopic analysis impact data resolution, with profound consequences for understanding past agricultural diversity.

## 1 Introduction

Over the past ten years, stable carbon (δ^13^C) and nitrogen (δ^15^N) isotope analyses of charred seed remains have become an increasingly important analytical tool used to understand ancient cultivation practices and agricultural systems (Styring et al., 2017a; Bogaard et al., 2019; Ferrio et al., 2020; Li et al., 2022; Nayak et al., 2022). Carbon and nitrogen stable isotope ratios measured from cereal grains, pulses, and other plant parts have revealed the ways in which people managed their plant resources while also providing insights into local environmental conditions that further shaped their plant management practices (Riehl et al., 2008; Riehl, 2012). Isotope analyses of ancient charred seeds have provided crucial insights into how small-scale watering practices, dedicated irrigation, and the application of manure to fields as fertilizer have shaped the cultivation systems developed by hunter-gatherer-cultivators, fisher-farmers, and full-time farmers alike (Ferrio et al., 2005; Fraser et al., 2011; Kanstrup et al., 2011; Bogaard et al., 2013; Stroud et al., 2021). In turn, these essential plant husbandry practices have revealed new insights into how people sustained and increased crop yields (Bogaard et al., 2013; Yang et al., 2022), took up and incorporated novel crops into extant agricultural systems (Li et al., 2022), promoted agricultural intensification and extensification (Styring et al., 2017a), and fostered social inequality and emerging social complexity (Bogaard et al., 2019).

These crucial perspectives on ancient agricultural practices rely on information derived from the isotope analysis of a relatively limited number of carbonized seeds recovered from archaeological deposits. Analyses of ancient carbonized seeds typically involve the selection of ca. 5 to 25 specimens, each representing the same taxon, recovered from a single archaeological depositional context (Fraser et al., 2011), followed by the aggregation and homogenization of these seeds to form a single pooled sample that is then measured for carbon and nitrogen isotope ratios (Nitsch et al., 2015; Styring et al., 2017b). The rationale underlying this approach is that any isotopic differences between single grains would be effectively averaged to obtain a more robust and representative stable isotope ratio that fully encompasses growing conditions represented by the plant remains recovered from that context (Nitsch et al., 2015; Styring et al., 2017b). The recommendation to aggregate specimens also reflects an analytical choice to ensure the isotope ratio mass spectrometer has an adequate amount of nitrogen (%N) to reliably obtain nitrogen isotope values (Vaiglova et al., 2023). However, ongoing developments in mass spectrometry and higher sensitivity in instrumentation have reduced the amount of nitrogen (%N) required in a carbonized sample, increasing the reliability of carbonized single grain measurements. Despite the increasing viability of conducting stable isotope measurements on single grains, relatively few studies have explored how different sampling strategies in stable isotope analysis impact data output and interpretation (Gavériaux et al., 2022; Gron et al., 2021; Larsson et al., 2019; Lightfoot and Stevens, 2012).

Stable isotope ratios measured from aggregated sets of charred grains effectively trace temporal shifts in agricultural practices that alter water availability and nitrogen levels in soils (Styring et al., 2017b; Bogaard et al., 2019; Riehl, 2020). However, the extraction of isotope values measured from individual grains, rather than a single isotope value that a pooled aggregate ‘bulk’ sample would provide, has considerable potential to more closely explore the diversity between and within ancient agricultural systems and better understand how cultivation practices changed over time in concert with or independently of broader shifts in environmental conditions, socio-cultural dynamics, and modes of economic production. As such, isotopic differences between individual carbonized grains found in archaeobotanical assemblages can identify variation between watering and manuring practices, which aggregate samples may obscure, masking temporal and spatial variation within past growing conditions (Lightfoot and Stevens, 2012; Larsson et al., 2019). Recent discussion centered on single-grain and aggregate sampling note that single-grain measurements are useful for understanding intra-context variability, while aggregate samples are more suitable for primary contexts where archaeobotanical assemblages represent a single years’ harvest (Vaiglova et al., 2023). A single-grain isotope value represents highly specific, spatially bounded anthropogenic and environmental inputs that impact a plants growth (Bogaard et al., 2007). In contrast, aggregate samples, in essence, create an isotope average (IA) composed of multiple charred grains, which, even if recovered from the same archaeological context, may have originated from different agricultural fields subjected to widely different cultivation practices or localized growing conditions (Vaiglova et al., 2023). Furthermore, the resulting aggregated IA value would likely erase isotopic variation caused by, for example, differences in water availability between annual harvests (Flohr et al., 2011), or manuring priorities (Styring et al., 2016b), collapsing spatial variation in growing conditions (Lightfoot and Stevens, 2012). Alternatively, an IA may obscure cases where there is, in actuality, very little carbon or nitrogen isotope variation across grains reflecting homogeneity in cultivation practices and growing conditions (Gavériaux et al., 2022). In addition, long term challenges in consistent reporting of results and scientific reproducibility can arise due to inter-aggregate sample dissimilarities in the numbers of grains (between 5 to 25) that comprise a single aggregate sample (Nitsch et al., 2015; Gron et al., 2021). The isotope value representing an aggregated sample is a mean value without a standard of deviation, standard of error, median, or variance, erasing measures of uncertainty before they can be assessed (Cowgill, 2015; Calin-Jageman and Cumming, 2019; Drennan, 2009; Shennan, 2006). In practice, these pitfalls might be overcome through analysis of large datasets composed of hundreds of aggregate samples that might reasonably record wide regional and temporal ranges of variation (e.g. Styring et al., 2017a), or avoided if depositional contexts representing a very short formation period are sampled based on the assumption that recovered archaeobotanical remains would represent a temporally constrained assemblage and thus represent a narrow range of growing conditions (Styring et al., 2016b).

Here, we investigate how carbon and nitrogen isotope values measured from single-grains or aggregated sample sets impact the average isotope values and variation represented in a cohesive dataset, and, as such, our interpretations of ancient plant management practices detectable though the stable isotopic record. We first analyze modern barley (*Hordeum vulgare* var. *nudum*) grown from an uncontrolled experimental plot, presenting the first analysis of intra-panicle variation of both carbon and nitrogen values from the same panicle. This expands the empirical references for single-grain isotope variation available to archaeologists. These results are then compared with the carbon and nitrogen composition of both charred and uncharred modern grains to understand how representative the isotope values of a single grain may be of its proximate growing conditions, as well as how those values might be transformed through charring. We expand our analysis by directly comparing isotope values measured from ancient single-grain and aggregate samples recovered from the same archaeological context in order to evaluate the variation represented in single-grain and aggregate samples. To this end, we measured carbon and nitrogen isotope ratios of charred single grains and aggregate samples consisting of multiple individual specimens of hulled barley (*Hordeum vulgare*) from late urban contexts at the Indus civilization site of Harappa, Pakistan (Periods 4/5, 1900-1700 BCE). We examine the results from the single-grain and aggregate samples to compare potential interpretations of each sample set, discussing the implications for understanding past agricultural organization and land use.

## 2 Isotopic variation in aggregate samples and single grains

Single-grain isotope analyses are increasingly used to make use of limited archaeobotanical material, or to understand variation between single archaeological contexts or sites (Riehl et al., 2008; Larsson et al., 2019; Vaiglova et al., 2020; Gron et al., 2021; Li et al., 2022). However, there remains little direct discussion of the use of single grains over aggregate samples in archaeological contexts (Vaiglova et al., 2023). Isotope data derived from archaeological grains provide invaluable information on anthropogenic and environmental conditions present during crop cultivation. The relationships between watering or manuring conditions and cultivar δ^13^C and δ^15^N values have been established through the isotopic analyses of grains collected from experimentally grown plots of wheat, barley, and other crops (Flohr et al., 2011; Fraser et al., 2011; Kanstrup et al., 2011; Wallace et al., 2013; Styring et al., 2016a). In general, the relationship between these inputs and isotope values and variation has been established using aggregate samples of modern grains from such experimental fields, and the interpretative framework developed by this work is applied to both single-grain and aggregate archaeological samples, there remains little direct discussion of the use of single grains over aggregate samples in archaeological contexts (i.e., Vaiglova et al., 2023).

### 2.1 Carbon variation in aggregate sample and single-grain experimental studies

To investigate the impact of irrigation and aridity on crop carbon isotope values, studies have relied on outdoor growing experiments that correlate broad categories of water availability with δ^13^C and carbon isotope discrimination (Δ^13^C) values in aggregate samples (Wallace et al., 2013; Styring et al., 2016a). Crop carbon stable isotope values are primarily influenced by photosynthetic pathway and water availability (Araus et al., 1997a; Cappers and Neef, 2012; Farquhar et al., 1989; Tieszen, 1991; Vogel, 1993), although field proximity to forests where light availability may be lower would also impact crop δ^13^C values (Van Der Merwe, 1982; van der Merwe and Medina, 1991; Bonafini et al., 2013). Internal stomatal conductance of CO_2_ also further modifies the carbon isotope discrimination of plants, with water-efficient flora typically exhibiting lower stomatal conductance and/or higher photosynthetic capacity that consequently impart an ^13^C-enrichment in plant tissues (Ma et al., 2020). This is reflected in the carbon isotopic composition of wheat and barley grown under the same conditions, with barley δ^13^C and Δ^13^C values approximately 1‰ higher than wheat (Flohr et al., 2019; Styring et al., 2016a; Wallace et al., 2013).

Modern aggregate samples show limited carbon isotope variation in crop plants grown in the same field, whereas more significant differences could exist between crops grown in different fields under similar watering conditions (Ferrio et al., 2005; Wallace et al., 2013; Jones et al., 2021). Research establishing the initial frameworks for assessing water availability drew on 168 samples, each consisting ∼25 to 50 aggregated grains of wheat and barley grown in arid and semi-arid fields in Spain and Syria were collected from a mix of rainfed and irrigated water regimes. Aggregated samples of barley harvested in Spain in 2007 and 2008 demonstrated overall low variation in mean Δ^13^C values. In unirrigated (n = 2) and moderately irrigated (n = 3) barley from a single year’s harvest, these differences were extremely limited (2008: 18.9 ± 0.2‰ vs. 18.7 ± 0.5‰), but fully irrigated fields (2007: n = 6, 2008 n = 3) exhibited higher mean Δ^13^C values as well as variation (2007: 18.8 ± 0.6‰, 2008: 19.6 ± 1.1‰) (Wallace et al., 2013). This higher variation in fully irrigated wheat mean Δ^13^C values in different years (2007: n = 5, 2008: n = 3) was not as pronounced (2007: 17.81 ± 0.81‰, 2008: 18.14 ± 0.56‰) (Wallace et al., 2013). Similar ranges of limited intra-field variation was also observed in aggregate samples (n_aggregate_ = 36, each of 50 grains) in plants subjected to three distinct regions with different watering in semi-arid fields in Morocco (Styring et al., 2016a). Two of the regions, Rainfed North and Rainfed South reflect the overall water availability and precipitation, with average carbon isotope values of −27.6‰ ± 0.5‰ (703mm) and −23.6‰ ± 0.6‰ (272mm) respectively. The irrigated and flood-cultivated Oasis crops exhibited a mean δ^13^C value of −26.3 ± 0.6‰ with similar watering conditions to Rainfed North. There is extremely limited overall variation between aggregate samples within each region of ± 0.6‰ (Styring et al., 2016a).

Notably, the Δ^13^C values of aggregate barley (n =10 grains) grown in unirrigated rainfed fields in Jordan showed greater carbon isotope differences between different year’s harvests from the same field (n = 3 per year, 2005–2006: 15.4 ± 0.1‰, 2006–2007: 16.5 ± 0.2‰, 2007–8: 17.3 ± 0.1‰) (Flohr et al., 2019). In addition to these temporal differences, Jones et al. (2021) documented spatial differences in Δ^13^C values of 2‰ between barley grown in different fields with similar watering conditions across Northwest India. In 124 aggregate samples (n_grain_ = 20–30 grains each) from uncontrolled flooded (n_sample_ = 101, 18.0 ± 1.5‰), sprinkler-irrigated (n_sample_ = 19, 16.8 ± 1.4‰), and rainfed fields (n_sample_ = 4, 17.2 ± 2.8‰), this study found that while water availability was the primary driver of carbon isotope variation, the variation between fields with the same watering conditions could be significant (Jones et al., 2021).

Single-grain isotope analysis has found limited intra-panicle and intra-field carbon isotope variation (Heaton et al., 2009). In two bread wheat (*Triticum aestivum* ssp. *vulgare*) plants grown in the same uncontrolled field in Nottingham (UK) (n_grain_ = 18), single-grain δ^13^C values ranged between –26.5 to –27.5‰. In corresponding aggregate samples from the same fields, each consisting of ∼300 uncarbonized grains from 6 panicles, yielded δ^13^C standard deviations between ± 0.37 to ± 0.83‰ (Heaton et al., 2009). These studies have found consistent limited ranges of δ^13^C and Δ^13^C variation found in aggregate and single-grain samples of grains grown in the same field and year (Araus et al., 1997b; Styring et al., 2016a; Flohr et al., 2019; Wallace et al., 2013; Araus et al., 2003). However, more notable variation in carbon isotope values is possible between inter-annual harvests from the same fields, as well as spatial variation independent of watering conditions (Wallace et al., 2013; Flohr et al., 2019; Jones et al., 2021). These patterns of carbon isotope variation indicate that information of watering practices may be lost in aggregate samples if grains originate from different fields or years.

### 2.2 Nitrogen variation in aggregate and single-grain experimental studies

Crop nitrogen isotopic composition is driven by plant uptake of bioavailable nitrogen through either nitrate (NO_3_^-^) or ammonium (NH_4_^+^) (Denk et al., 2017), both of which are affected by manuring and soil moisture (Craine et al., 2015; Szpak et al., 2017). Grain δ^15^N values reflect nutrient availability during grain-filling, and practices such as manuring enrich soil ^15^N through the direct addition of ammonia to soils, driving floral nitrogen isotope values up, sometimes considerably (up to 10‰) (Fraser et al., 2011; Kanstrup et al., 2011; Szpak, 2014; Craine et al., 2015). In addition to marked ^15^N-enrichment in grain values visible in cultivars grown under manured conditions (Bogaard et al., 2007; Chadwick et al., 2000; Aguilera et al., 2008; Fraser et al., 2013; Styring et al., 2016a), there can be differences in the amplitude of nitrogen isotopic variation for grains from manured plots compared to those from unmanured plots, but this appears to be dependent in part on regional environmental conditions. In controlled experimental plots located in temperate environments in Hertfordshire (UK), aggregate unmanured and manured samples yield widely different δ^15^N values but similar ranges of variation (δ^15^N_manured_ = 1.9 ± 1.4‰, range = 0.6 to 4.0‰; δ^15^N_manured_ 7.4 ± 1.1‰) (Bogaard et al., 2007; Fraser et al., 2013). In contrast, aggregate samples from Morocco found dramatic differences in nitrogen isotope values between manured and unmanured fields, but low nitrogen isotope variation in samples from unmanured fields (Rainfed North δ^15^N_mean_ 0.8 ± 0.6‰) and irrigated manured fields (Oasis δ^15^N_mean_ 14.0 ± 1.0‰), but also wide variation in rainfed manured fields (Rainfed South δ^15^N_mean_ = 7.8 ± 2.1‰).

Analyses of single-grains also found slightly greater variation in manured versus unmanured grains (Bogaard et al., 2007; Larsson et al., 2019). In plants from the same fields, single-grain intra-panicle variation in unmanured wheat δ^15^N values was limited to ± 0.1‰ (range = –0.5 to 0.5‰) while manured grains varied by ± 1.3‰ (range = 5 to 7.5‰) (Bogaard et al., 2007). A similar pattern was identified in manured and unmanured experimental plots in Borgeby, Sweden where intra-panicle variation in δ^15^N values measured from the single grains of unmanured 2-row hulled barley (*Hordeum vulgare* ssp. *distichon*) averaged 5.4‰ ± 0.6‰ (range = 4.1 to 6.2‰.), but in manured grains 8.9 ± 1.7‰ (range = 6.4 to 11.8‰) (Larsson et al., 2019). The variation observed between individual grains from the same fields (ca. 1–2‰), and the relatively high variation present between some aggregate sample wheat and barley δ^15^N mean values, suggests that important information on fertilization practices may be heavily obscured in these pooled samples.

### 2.3 The impact of charring on charred seed isotope values

Archaeobotanical assemblages are in most cases made up of plant material preserved through charring, and extensive research has been conducted on the impacts of carbonization on grain morphology and taphonomy (Boardman and Jones, 1990; Charles et al., 2015; Hillman et al., 1993; Pearsall, 2015; van der Veen, 2007). The increasing use of stable isotope analysis on charred grains has extended this work to investigating the charring impacts on the original carbon and nitrogen isotope values through experimental research on modern grains (DeNiro and Hastorf, 1985; Aguilera et al., 2008; Kanstrup et al., 2012; Fraser et al., 2013; Nitsch et al., 2015; Stroud et al., 2023a). However, these charring studies have focused their analysis on randomly selected grains from multiple plants in a field, which are carbonized together and aggregated into a single sample, then compared with a corresponding sample of randomly selected and aggregate uncharred grains. This has meant that no analysis has compared the intra-panicle variation of charred and uncharred single grains or explored any potential differences between single-grain isotope values resulting from carbonization.

Grains exposed to temperatures above 200°C undergo Maillard reactions that volatilize carbon (C-) and nitrogen (N-) containing compounds as seed starches convert to dextrin (Pazola and Cieslak, 1979; Styring et al., 2013). Grain %C and %N rise by ca. 20% and 2.5%, respectively, while grains lose 20 to 40% of their mass, proportional to temperature (Czimczik et al., 2002; Braadbaart et al., 2004; Kanstrup et al., 2012). Between 230 to 300°C, these reactions leads to some loss of ^14^N and increase in δ^15^N values (Bogaard et al., 2007; Kanstrup et al., 2012; Styring et al., 2013), while δ^13^C values remain largely unchanged (Aguilera et al., 2008; Fraser et al., 2011; Nitsch et al., 2015). At temperatures above 300°C, gross deformation in grain morphology occurs, with mass loss over 50% (Boardman and Jones, 1990; Braadbaart et al., 2004; Charles et al., 2015). This is accompanied by significantly deviations from *in vivo* isotope values, with observed shifts of ≥ 1‰ in δ^13^C and ≥ 2‰ in δ^15^N values (Czimczik et al., 2002; Kanstrup et al., 2012; Fraser et al., 2013; Stroud et al., 2023a).

Many studies of charring impacts have compared the mean carbon and nitrogen values between aggregate charred and uncharred samples. In grains charred between 200°C and 300°C a mean increase of +0.6‰ in the δ^15^N values of 5 samples out of 15 grains was found when compared with uncharred aggregate samples (Kanstrup et al., 2011, 2012). Fraser et al. (2013) charred aggregate samples of 25 to 50 grains from Syria, Germany, and the United Kingdom at 230°C, which when compared with uncharred aggregate samples from the same fields, reported higher charred δ^15^N values up to +0.8‰ but no notable differences in δ^13^C values (Bogaard et al., 2007; Fraser et al., 2011).

Other research has assessed charring effects on grain isotope values using statistical correlations or linear regression models to predict isotopic offsets for archaeobotanical grain isotope samples. Aguilera et al. (2008) compared samples of five aggregate wheat or barley grains charred at 250°C with uncharred samples, testing correlations between uncharred and charred samples, suggesting a δ^15^N offset of ca. 0.68‰ to compensate for charring impacts (Aguilera et al., 2008). Subsequent work proposed offsets derived from modern charring experimental data and multiple linear regression models incorporating, time and temperature to predict impacts on archaeological δ^13^C and δ^15^N values (Nitsch et al., 2015; Stroud et al., 2023a). In unmanured samples of eight taxa, including bread wheat (*Triticum aestivum)* and hulled barley (*Hordeum vulgare*), samples of 10 grains were fired between 230 to 300°C between 4 to 24 hours and compared with uncharred samples. For every 15°C above 200°C, δ^13^C vales increased by 0.05‰, and δ^15^N by 0.12‰, whereas every four hours of charring resulted in a 0.016‰ increase δ^13^C, and 0.04‰ for δ^15^N values (Nitsch et al., 2015; Stroud et al., 2023a). From these results, Nitsch et al. (2015) predicted overall isotopic offsets of 0.31‰ for δ^15^N values, and 0.11‰ in δ^13^C values of grains carbonized between 230 to 260°C. Stroud et al. (2023) building upon this data argued for offsets of 0.32‰ in δ^15^N, and 0.16‰ in δ^13^C to be subtracted from archaeological wheat and barley grain isotope values (Stroud et al., 2023b).

### 2.4 Aggregate sampling strategies in archaeobotanical material

Experimental studies of modern material have repeatedly shown both the viability of correlating charred grain stable isotope values to cultivation practices and environmental conditions. However, most experimental research has assessed isotopic variability between fields or the charring impacts by aggregate samples of modern grains grown under known conditions. Archaeobotanical grains complicate the use of aggregate samples as they cannot be assumed to derive from a single field or set of growing conditions.

The clearest argument making the case for aggregating archaeological grains derives from work applying a multiple linear regression model calculating potential charring offsets (Nitsch et al. 2015). These studies used this model to define the range of carbon and nitrogen isotope variation within a single modern growing context (e.g., a field), and define adequate sample sizes for archaeobotanical stable isotope analysis (Nitsch et al., 2015; Stroud et al., 2023a). The analysis, based on seventy aggregate samples of ten grains from eight taxa from unmanured fields, calculated a residual standard error (SE) of ca. 0.25‰ in δ^13^C and 0.5‰ in δ^15^N. From this calculation, this work argued that within a 95% confidence interval, any expected variation in grain δ^13^C and δ^15^N values from a single field would be approximately ± 0.5‰ in δ^13^C, and ± 1.0‰ in δ^15^N (1.96 x SE) (Nitsch et al., 2015; Stroud et al., 2023a). Nitsch et al. (2015) hypothesized that decreasing the number of grains homogenized together within a single aggregate sample would in turn increase the standard error, increasing the variation and uncertainty of a single measurement, thereby rendering individual grain isotope values too variable to be interpreted (Nitsch et al., 2015; Stroud et al., 2023a). From these findings, drawing on aggregate samples from modern growing contexts, Nitsch et al. (2015) called for the inclusion of ten archaeological grains from a single archaeological context to create a single aggregated sample. This is intended to balance the preservation of archaeobotanical assemblages and as a means of increasing the sample size represented in a single stable isotope measurement from an archaeological context (Nitsch et al., 2015; Stroud et al., 2023b). However, the leap from known modern growing conditions to ultimately uncertain archaeological data, renders the assessment of single grain isotope analysis necessary. Despite this, no formal comparisons of single grain isotope values or aggregate sample values have yet been carried out.

## 3 Materials and Methods

### 3.1 Barley grains from an experimental plot at Steinzeitpark Dithmarschen

We performed δ^13^C and δ^15^N isotope analyses of single grains from the same panicle to better define the range of intra-panicle isotopic variation for unmanured crops growing in a well-watered C_3_ environment and assess the impacts from charring in that variation. Four panicles of naked barley (*Hordeum vulgare* var *nudum*) were selected from legacy collections generated by growing experiments conducted by Institute of Prehistoric and Protohistoric Archaeology, University of Kiel at Steinzeitpark Dithmarschen, Albersdorf, Germany (Figure 1). The ca. 5m x 5m plot lies on the grounds of the Steinzeitpark Dithmarschen, with sandy clay soils that have not been manured or received fertilizer since at least 2005, when the park was founded (Burbaum et al., 2019; Beuker, 2020). The barley was sown by hand in April 2017, left to grow with no intervention until harvest, receiving no manure or additional water beyond environmental precipitation of ca. 315mm (Deutscher Wetterdienst, 2024), with mature ripe plants harvested in August 2017. Each panicle was assigned a letter (A, B, D, E). Grains on each panicle were sequentially numbered in ascending order, from the base to the apex, recording their relative position (Figure 2). All grains from two panicles, D (n = 33) and E (n = 30), were individually measured for carbon and nitrogen isotopes in order to assess intra-panicle isotopic variation within and between single inflorescences. These uncarbonized grains were ground individually into powder using an agate mortar and pestle and then weighed (2.5mg) into tin capsules for carbon and nitrogen isotope ratio mass spectrometry.

**Figure 1.**
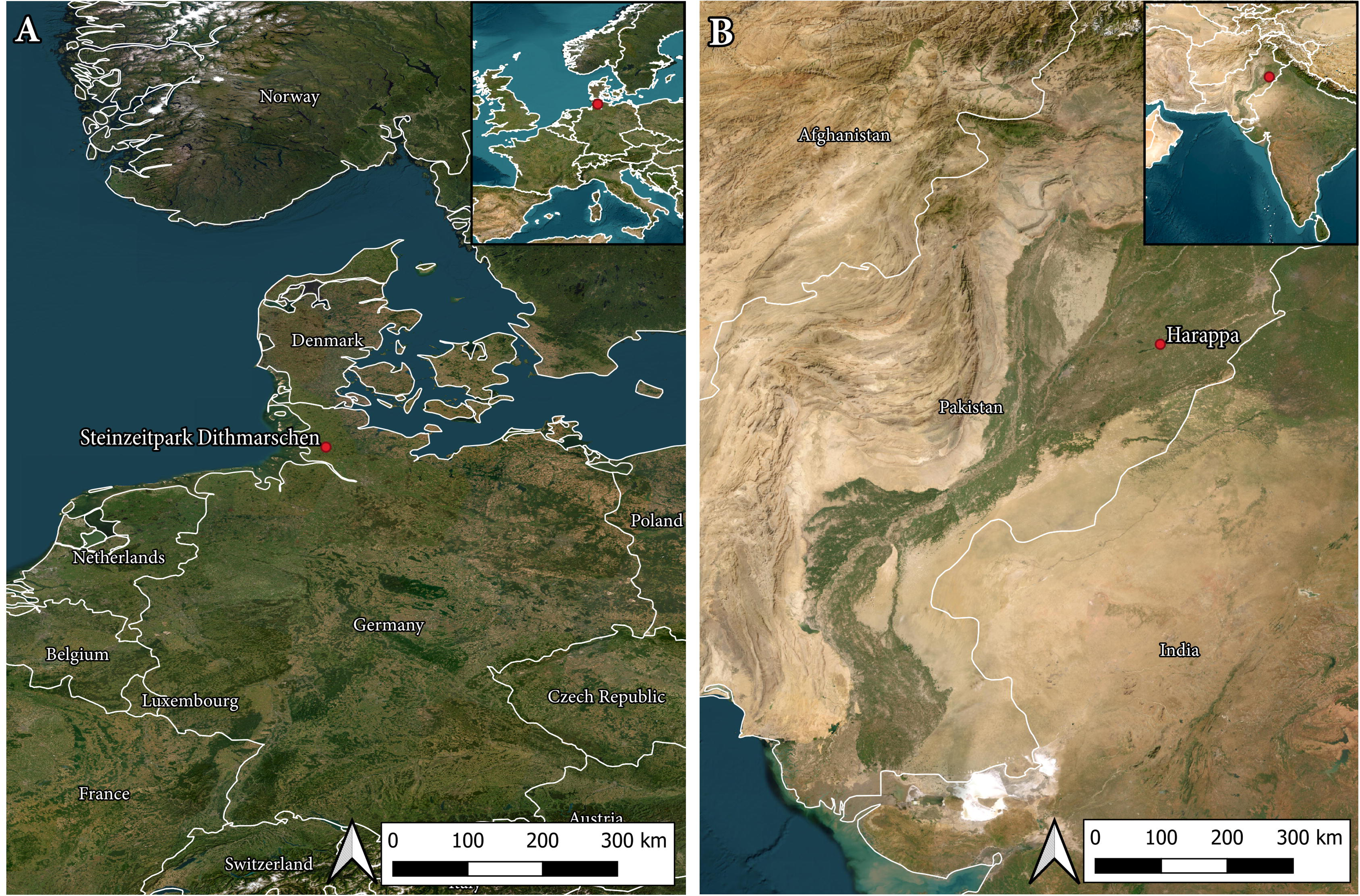
Map showing locations of Steinzeitpark Dithmarschen and Harappa.

**Figure 2.**
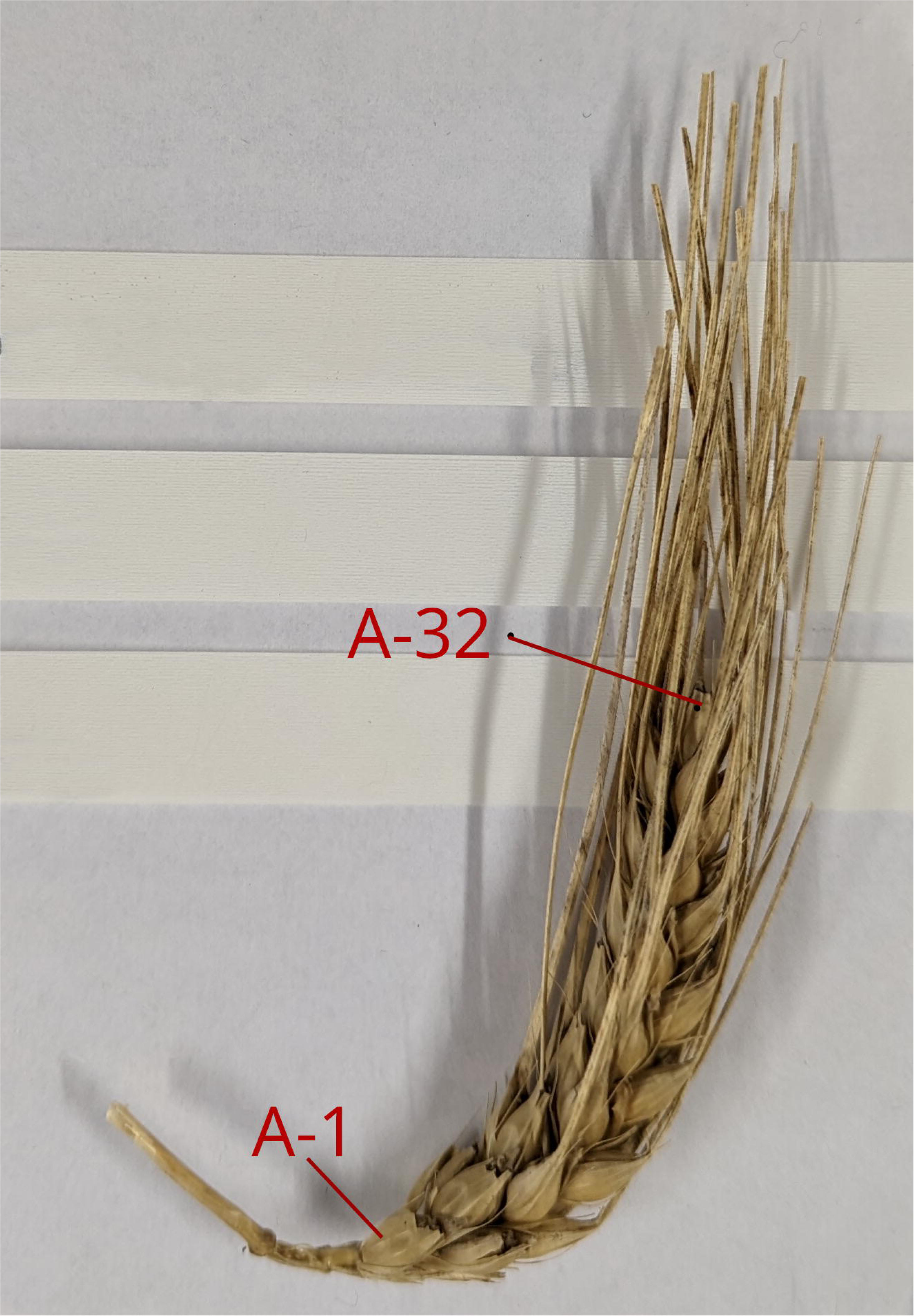
Example grain numbering of *Hordeum vulgare* var *nudum* Panicle A from Steinzeitpark Dithmarschen.

Plant remains recovered from archaeological contexts are typically preserved through exposure to fire (Gallagher, 2015). Studies of charring impacts on grain stable isotope values using aggregate samples have found minimal increases to grains charred between 230 to 300°C (Fraser et al., 2013; Vaiglova et al., 2014). To assess how charring potentially impacts the range and of intra-panicle isotopic variation, we measured individual grains subjected to a range of charring conditions. Four unaltered grains each from panicles A and B were selected for mass spectrometry and set aside. Additional 26 grains from panicles A and B were individually carbonized in both anoxic and oxidizing conditions at 230°C (n=8), 300 (n = 16), and 400°C (n = 4) in a muffle furnace (Table 1, Supplemental Information 1). Grains carbonized in anoxic conditions were individually wrapped in aluminum foil, with each foil packet then placed together in an enclosed ceramic crucible. Grains carbonized in oxidizing conditions were placed individually in an open ceramic crucible individually. All uncarbonized and carbonized grains were ground individually into powder using an agate mortar and pestle and then weighed (2.5mg) into tin capsules for carbon and nitrogen isotope ratio mass spectrometry.

**Table 1.** Summary statistics of δ^13^C and δ^15^N values for Steinzeitpark Dithmarschen Barley. Panicle A (Uncarbonized n = 4, 230°C n = 6), Panicle B (Uncarbonized n = 4; 230°C, n = 8; 300°C, n = 8), Panicle D (Uncarbonized n = 33), Panicle E (Uncarbonized n = 24).

| | | *n* | $\delta^{13}\text{C}$<br>*Mean* | $\delta^{13}\text{C}$<br>*SD* | $\delta^{13}\text{C}$<br>*Min* | $\delta^{13}\text{C}$<br>*Max* | $\delta^{15}\text{N}$<br>*Mean* | $\delta^{15}\text{N}$<br>*SD* | $\delta^{15}\text{N}$<br>*Min* | $\delta^{15}\text{N}$<br>*Max* |
| --- | --- | --- | --- | --- | --- | --- | --- | --- | --- | --- |
| **Panicle A** | Uncarbonized | 4 | -28.6 | 0.2 | -28.8 | -28.4 | 1.2 | 0.1 | 1.2 | 1.3 |
|  | 230°C | 6 | -28.5 | 0.3 | -28.9 | -28.1 | 1.4 | 0.6 | 0.2 | 2.0 |
|  | All Panicle A* | 10 | -28.5 | 0.3 | -28.8 | -28.1 | 1.3 | 0.4 | 0.2 | 2.0 |
|  | 400°C *not included | 2 | -26.0 | 0.8 | -26.6 | -25.4 | 4.0 | 1.2 | 3.1 | 4.8 |
| **Panicle B** | Uncarbonized | 4 | -29.1 | 0.3 | -29.5 | -28.8 | 1.0 | 0.4 | 0.5 | 1.3 |
|  | 230°C | 8 | -29.0 | 0.2 | -29.4 | -28.6 | 1.2 | 0.2 | 1.0 | 1.4 |
|  | 300°C | 8 | -29.0 | 0.4 | -29.7 | -28.5 | 1.5 | 0.2 | 1.2 | 1.8 |
|  | All Panicle B | 20 | -29.0 | 0.3 | -29.7 | -28.5 | 1.3 | 0.3 | 1.2 | 1.8 |
| **Panicle D** | Uncarbonized | 33 | -28.9 | 0.2 | -29.5 | -28.5 | 0.7 | 0.3 | -0.6 | 1.2 |
| **Panicle E** | Uncarbonized | 30 | -28.9 | 0.2 | -29.4 | -28.3 | -0.2 | 0.4 | -1.2 | 0.4 |
| **All Panicles** | Uncarbonized | 71 | -28.9 | 0.3 | -29.5 | -28.3 | 0.4 | 0.6 | -1.2 | 1.3 |
| **All Panicles** | All Carbonized | 24 | -28.6 | 0.4 | -29.7 | -28.1 | 1.3 | 0.4 | 0.2 | 2.0 |

### 3.2 Archaeological grains from a single context at Harappa, Trench 42

We expand our exploration of single-grain isotope analysis by examining the degree to which carbon and nitrogen stable isotope ratios derived from an aggregated pool of single grains encompass the range of isotopic variation exhibited by those single grains. This is done through the stable isotope analysis of aggregated and single-grain sample sets all obtained from the same primary archaeological context. Charred samples of hulled barley (Hordeum vulgare) were collected from a burned storage bin made of wattle and daub discovered by the Harappa Archaeological Project (HARP) in the eroded upper levels of Trench 42, on Mound AB at the Indus site of Harappa, Pakistan (Figure 1, ca. 1900-1700 BCE) (Meadow and Kenoyer, 2001). This charred bin and associated wash layers contained thousands of charred barley grains (Meadow and Kenoyer, 2001). Cemetery H ceramics associated with the context and two samples of charred barley that have been radiocarbon dated confirm that these samples date to Periods 4/5 ca. 1900-1700 BCE (Meadow and Kenoyer, 2001).

Covering over 150 ha in size, Harappa was a large urban center that may have supported a population of over ca. 40,000 people (Kenoyer, 2008; Kenoyer and Meadow, 2016). Harappa was linked with other Indus cities by trade (Kenoyer, 1997) and a regionally integrated material culture that included similar ceramic traditions and systems of weights (Possehl, 2002), set within a mosaic of dense rural settlements spanning a wide range of environmental conditions (Weber et al., 2010a; Petrie and Bates, 2017). Harappa is located adjacent to a paleochannel of the Ravi River and today receives ca. 250mm annual precipitation with ca. 90mm falling between November to April, and ca. 160mm between May to October (Pendall and Amundson 1990). Like other Indus sites, Harappa agricultural systems were structured by the South Asian monsoon (Bates, 2022; Clift and d’Alpoim Guedes, 2021; Weber, 2003; Fuller, 2006). Traditional farming practices for the region take advantage of seasonal precipitation peaks that support two cropping seasons: the summer (*kharif*) season when crops such as millets and grams are grown during the June to October monsoonal season, and the winter (*rabi*) seasons when barley, wheat, and lentils are grown between November to April (Cappers et al., 2016; Miller, 2006). Harappan agriculture likely followed a similar cropping system, suggested by the high ubiquity, counts, and weight of charred barley seeds, followed by winter-grown wheat and lentils, seeds found in the archaeobotanical assemblage (James et al., 2024; Weber, 2003; Weber et al., 2010a).

Large-grained domesticated barley, the primary crop cultivated throughout the duration of occupation at Harappa, was intensively exploited (James et al., 2024), although millets and grams increased in importance by 2,600 BCE (Weber et al., 2010a, 2010b). Floodplain cultivation or some form of localized irrigation potentially provided barley fields with an additional water source required to meet barley watering requirement–(Miller, 2006, 2015; You, 2019). Crop production at Harappa may have been further enhanced through the use of zebu cattle and water buffalo (Meadow, 1996; Patel and Meadow, 2017), which served as a source of manure (Lancelotti, 2018; James et al., 2024), as well as traction, suggested by pathologies on their skeletal extremities (Miller, 2004) and terracotta models of cattle pulling carts and plows (Kenoyer, 2004). Secondary products formed a substantial component of Harappa agricultural systems, with cattle, sheep, and goats also husbanded for their meat and milk (Meadow, 1996; Miller, 2003; Patel and Meadow, 2017).

### 3.3 Archaeological Materials: Trench 42 Barley

A total of 46 carbonized grains were selected from the Trench 42 burnt storage feature. Grains were randomly selected and placed into two sample groups, S and A, representing single grains (S) each individually analyzed for carbon and nitrogen isotope ratios and aggregated samples (A) representing multiple seeds mechanically homogenized together for subsequent isotope analyses. To assess if the range of isotopic variation measured from randomly selected single grains (S1) matched a paired sample of aggregated seeds (A1), twenty grains were collected from the Trench 42 barley. Ten (S1; grain n = 10) of these grains were each individually ground into a fine powder using an agate mortar with ∼2.5mg of the resulting powder weighed into a tin capsule for single-grain measurements. The remaining ten grains (total number of grains in sample A1, grain n = 10) were collectively homogenized into a single aggregate sample A1 (total number of grains in sample A1, n = 10). The powder from A1 was measured six times, with weighed into tin capsules (A1; n = 6).

The carbon and nitrogen isotope ratios and range of isotopic variation exhibited by an aggregated sample representing multiple barley seeds was then compared the isotopic composition of single grains also used to create the aggregate sample sets. Twenty-six single grains (S2, n=26) were individually homogenized with ∼2.5mg of each grains powder separately weighed into tin capsules for single-grain stable isotope measurements. The remaining powders from the S1 individual grains were retained and homogenized together into the aggregate sample A2 (total number of grains in sample, n=26). The powder from A2 was measured six times, with each sample weighed into tin capsules (A2; n = 6).

### 3.4 Mass spectrometry

Carbon and nitrogen isotope analyses were undertaken at the Archaeology Stable Isotope Laboratory (ASIL), Institute for Prehistoric and Protohistoric Archaeology, University of Kiel. Samples were measured using an isoprime visION continuous flow isotope ratio mass spectrometer coupled to a vario PYRO cube elemental analyzer (Elementar Analysesysteme GmbH, Langenselbold, Germany). Stable carbon and nitrogen isotopic compositions were calibrated relative to the VPDB (δ^13^C) and AIR (δ^15^N) scales using the glutamic acid standards USGS40 (δ^13^C −26.39 ± 0.04‰, δ^15^N −4.52 ± 0.06‰) and USGS41a (δ^13^C 36.55 ± 0.08‰, δ^15^N 47.55 ± 0.15‰) (Qi et al. 2003; Qi et al, 2016). Samples were measured in 14 analytical runs. Measurement uncertainty was monitored using internal standards wheat and millet. The isotopic compositions reported here for internal standards represent long term averages calibrated to VPDB and AIR with USGS40 and USGS41a (Supplementary Material 1). Precision (u(R_w_)) was determined to be ± 0.07‰ for δ^13^C and ± 0.16‰ for δ^15^N on the basis of repeated measurements of calibration standards, check standards, and sample replicates. Accuracy (u_(bias)_) was determined to be ± 0.12‰ for δ^13^C and ± 0.13‰ for δ^15^N on the basis of the difference between the observed and known δ values of the check standards and the long-term standard deviations of these check standards. The total analytical uncertainty was determined to be ± 0.19‰ for δ^13^C and ± 0.11 for δ^15^N (after Szpak et al., 2017).

Carbon isotopes in archaeological grains are most often expressed in analysis as carbon isotope discrimination (Δ^13^C) to accommodate for differences in past and modern atmospheric carbon dioxide (CO_2_) concentrations and atmospheric δ^13^C (Farquhar et al., 1989; Araus et al., 1997a; Wallace et al., 2015; Rosen et al., 2019). Here, all δ^13^C values measured from archaeological seeds are presented as Δ^13^C calculated using an estimated δ^13^C_air_ of 6.4‰ (Farquhar et al., 1989; Francey et al., 1999; Ferrio et al., 2005). This calculation indicates the carbon isotope values of the archaeological seed in relation to the source of atmospheric CO_2_ i.e., Δ reflects the difference in δ^13^C values between the air and plant tissue (Farquhar et al., 1989).

## 4 Results

### 4.1 Intra-panicle carbon and nitrogen variation and composition of uncarbonized and carbonized modern barley from a C_3_ environment

Uncarbonized grains (n = 71) from the four panicles of modern, experimentally grown barley exhibited low variation in δ^13^C averaging –28.9 ± 0.3‰ (range = –29.5 to –28.5‰) (Table 1, Figure 3). The number of grains sampled per panicle influenced the isotopic variation expressed within each panicle. Panicle A (n = 4) yielded on average a δ^13^C value of –28.7 ± 0.2‰ (range = –28.8 to –28.4‰), Panicle B –29.1 ± 0.3‰ (n = 4, range = –29.5 to –28.8‰), Panicle D (n = 33) –28.9 ± 0.2‰ (n = 33, range = –29.5 to –28.5‰), and Panicle E –28.9 ± 0.2‰ (n = 30, range = –29.4 to –28.3‰).

**Figure 3.**
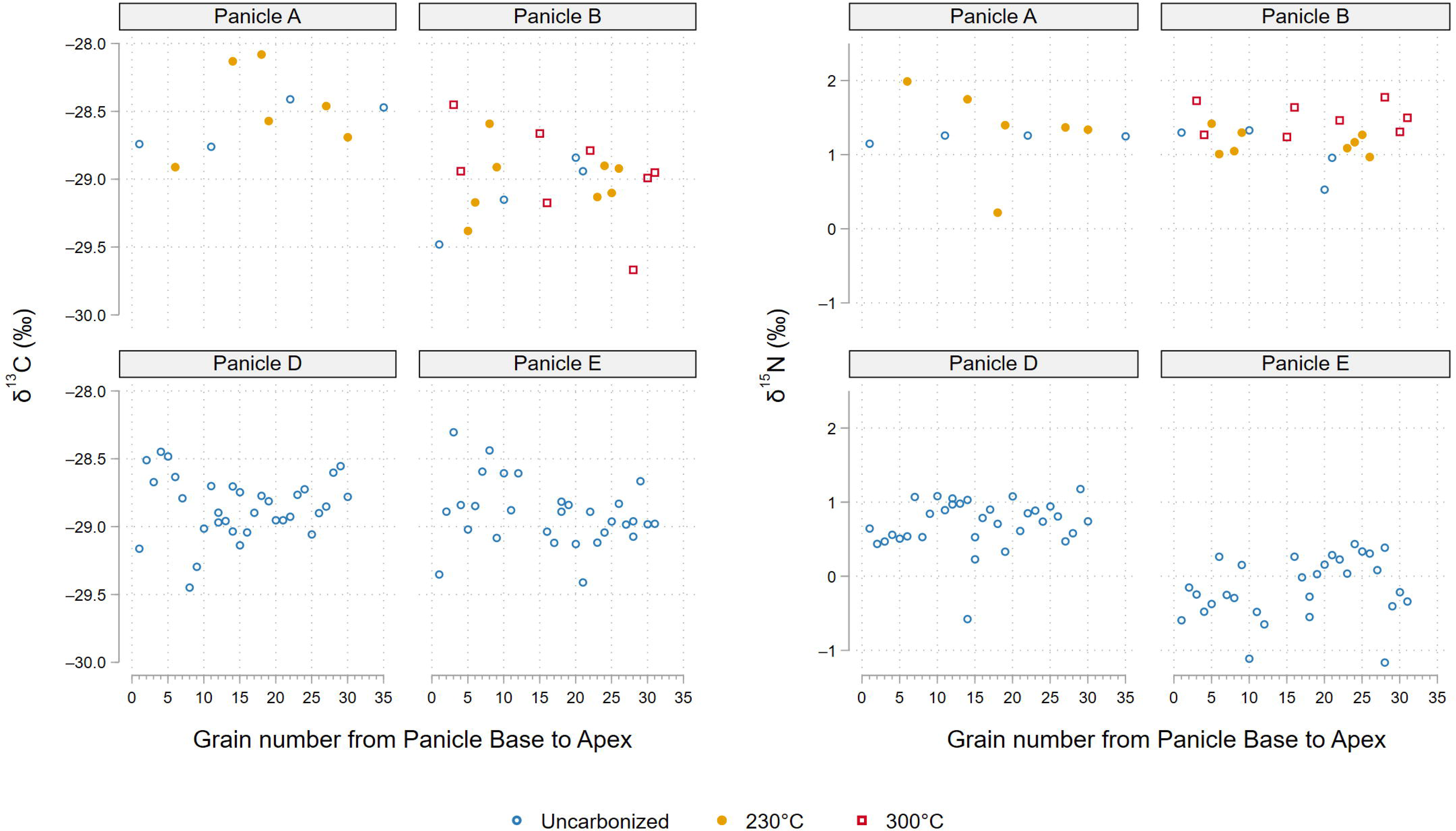
Steinzeitpark Dithmarschen single grain δ^13^C and δ^15^N values by panicle. Grains numbered from panicle base to apex.

For nitrogen isotopes, uncarbonized grains yielded an average δ^15^N value of 0.4 ± 0.6‰ (range = –1.2 to 1.3‰). Panicle A exhibited a mean *δ*^15^N value of 1.2 ± 0.1‰ (range = 1.2 to 1.3‰), Panicle B 1.0 ± 0.3‰ (range = 0.6 to 1.0‰), Panicle D 0.7 ± 0.3‰ (range = –0.6 to +1.2‰), and Panicle E –0.2 ± 0.4‰ (range = –1.2 to 0.4‰). No differences in δ^13^C or δ^15^N values based on the position of each grain on its respective panicle was observed (Figure 3).

Panicle A grains carbonized at 230°C (n = 6) averaging –28.5 ± 0.3‰ in δ^13^C (range = – 28.9‰ to –28.1‰) were similar in their isotopic composition as uncarbonized grains (Table 1, Figure). Panicle B grains carbonized at 230°C (n = 8) also yielded similar carbon isotope values as uncarbonized grains from the same panicle, with an average of –29.0 ± 0.1‰ (–29.4 to –28.6‰). Panicle B grains carbonized at 300°C (n = 8) also yielded similar carbon isotope values as uncarbonized grains averaging –29.0 ± 0.1‰ in δ^13^C (range = –29.7 to –28.5‰). In nitrogen isotopes, grains from Panicle B carbonized at 230°C (n = 8), seeds averaged 1.2 ± 0.2‰ in *δ*^15^N (range =1.0 to 1.4‰), similar to uncharred grains from the same panicle. Panicle B seeds carbonized at 300°C (n = 8) exhibited on average 0.5‰ in δ^15^N relative to uncarbonized seeds from the same panicle with an average of 1.5 ± 0.2‰ (range = 1.2 to 1.8‰).

Grains carbonized at 400°C (n = 4) from Panicle A exhibited strongly different carbon and nitrogen isotope values with two specimens ashed during the firing process. The grains (n = 2) yielded an average δ^13^C of –26.0 ± 0.6‰ (range = –26.6 to –25.4‰) and an average δ^15^N of 4.0 ± 1.2‰ (range = 3.1 to 4.8‰).

Steinzeitpark barley from Panicles A and B carbonized at 230°C and 300°C showed overlapping ranges of carbon and nitrogen isotope variation with the corresponding uncarbonized grains from those panicles. Inter-panicle differences in δ^13^C were low, but more pronounced in δ^15^N regardless of charring condition; this is particularly clear in the diverging nitrogen values of Panicle E (Table 1, Figure 4).

**Figure 4.**
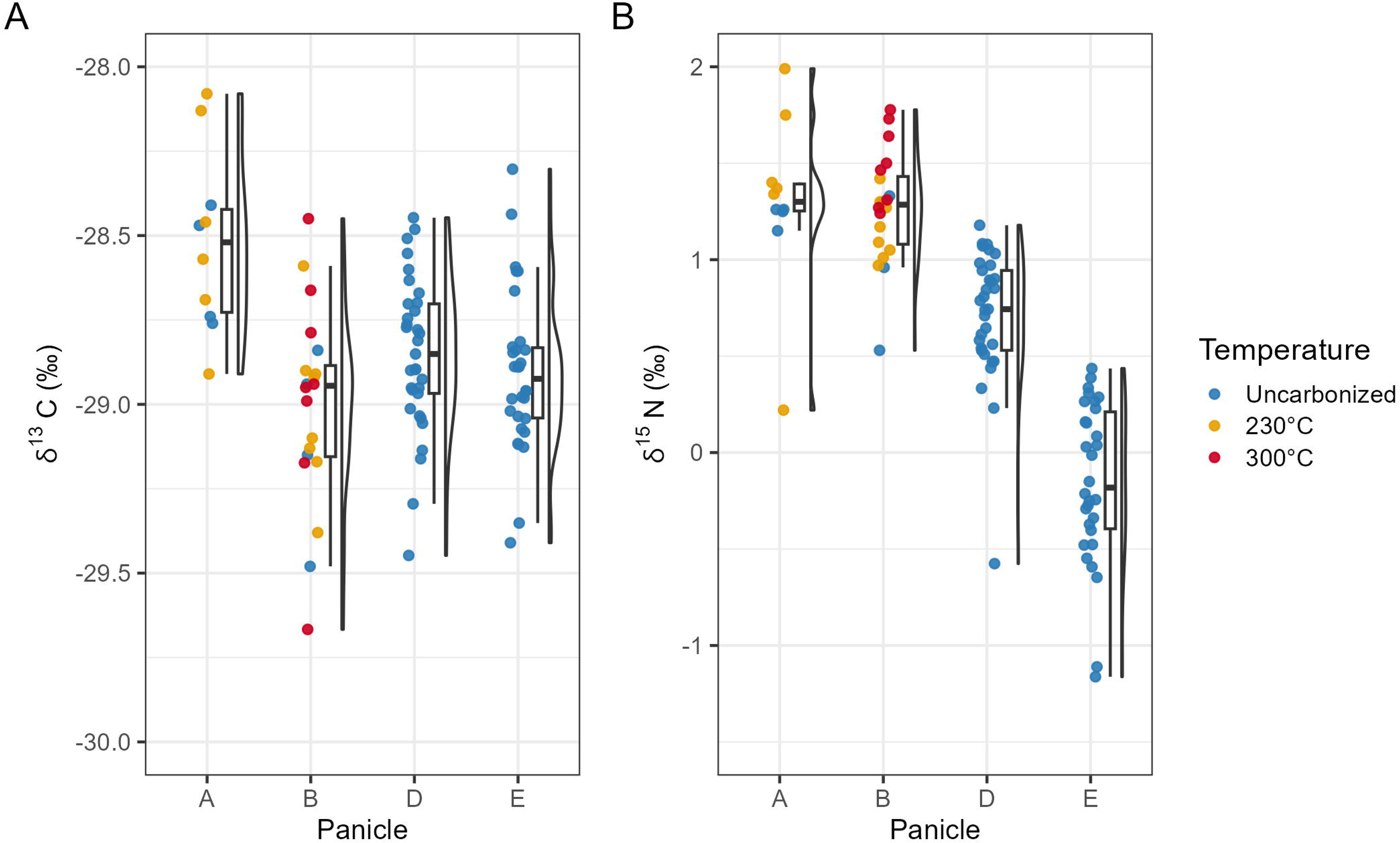
Boxplot of Steinzeitpark Dithmarschen barley isotope data. δ^13^C and δ^15^N values by panicle and temperature**. Panicle A** (Uncarbonized n = 4, 230°C n = 6)**, Panicle B** (Uncarbonized n = 4, 230°C n = 8, 300°C = 8), **Panicle D** (Uncarbonized n = 33)**, Panicle E** (Uncarbonized n = 24).

### 4.2 Carbon discrimination (Δ^13^C) and nitrogen (δ^15^N) variation in Harappa aggregate and single grain samples

Analysis of single grain and aggregate measurements of barley from Harappa Trench 42 found extremely wide variation in single-grain isotope values but notable overlap with isotope values measured from aggregate samples (Table 2, Figure 5). Across the 46 individually sampled grains, one sample failed due to low %N (Supplemental Information 1). Single grain measurements (n = 35) averaged 18.3 ± 0.8‰ in Δ^13^C (range = 16.7 to 20‰) and 5.2 ± 0.6‰ in δ^15^N. Two grains yielded outstandingly high δ^15^N values of 16 and 16.4‰, respectively; removal of these outliers results for an average δ^15^N value of 4.6 ± 1.9‰ for single grains (range = 0.6 to 8.1‰) (Table 1, Figure 5). S1 grains show average Δ^13^C values of 18.3 ± 0.8‰ (n = 9; range =16.7 to 19.4‰) and δ^15^N of 6.8 ± 4‰.The corresponding aggregate sample A1 (grain n =10) show a mean Δ^13^C value of 18.8 ± 0.1‰, and δ^15^N of 4.5 ±0.2‰, enriched on average 2.3‰ in ^15^N relative to the S1 seeds. The average carbon discrimination values of S2 and A2 are effectively identical, with slight differences between their average nitrogen isotope values. S2 (n = 26) grains average Δ^13^C value is 19.0 ± 0.7‰ (range =17.4 to 20‰) and S2 mean δ^15^N values are 4.6 ± 2.9‰ (range = 0.6 to 16.4‰). The corresponding aggregate sample A2 (grain n = 26) yielded a mean Δ^13^C of 19.0 ± 0.03‰ and 4.3 ± 0.2‰ in δ^15^N. There are no significant differences between the single grain or aggregate samples in either Δ^13^C (Kruskal-Wallis: H(1, 43) = 7.61, p = 0.643) or δ^15^N values (Kruskal-Wallis: H(1, 43) = 3.59, p = 0.742) (Table 2, Figure 6).

**Figure 5.**
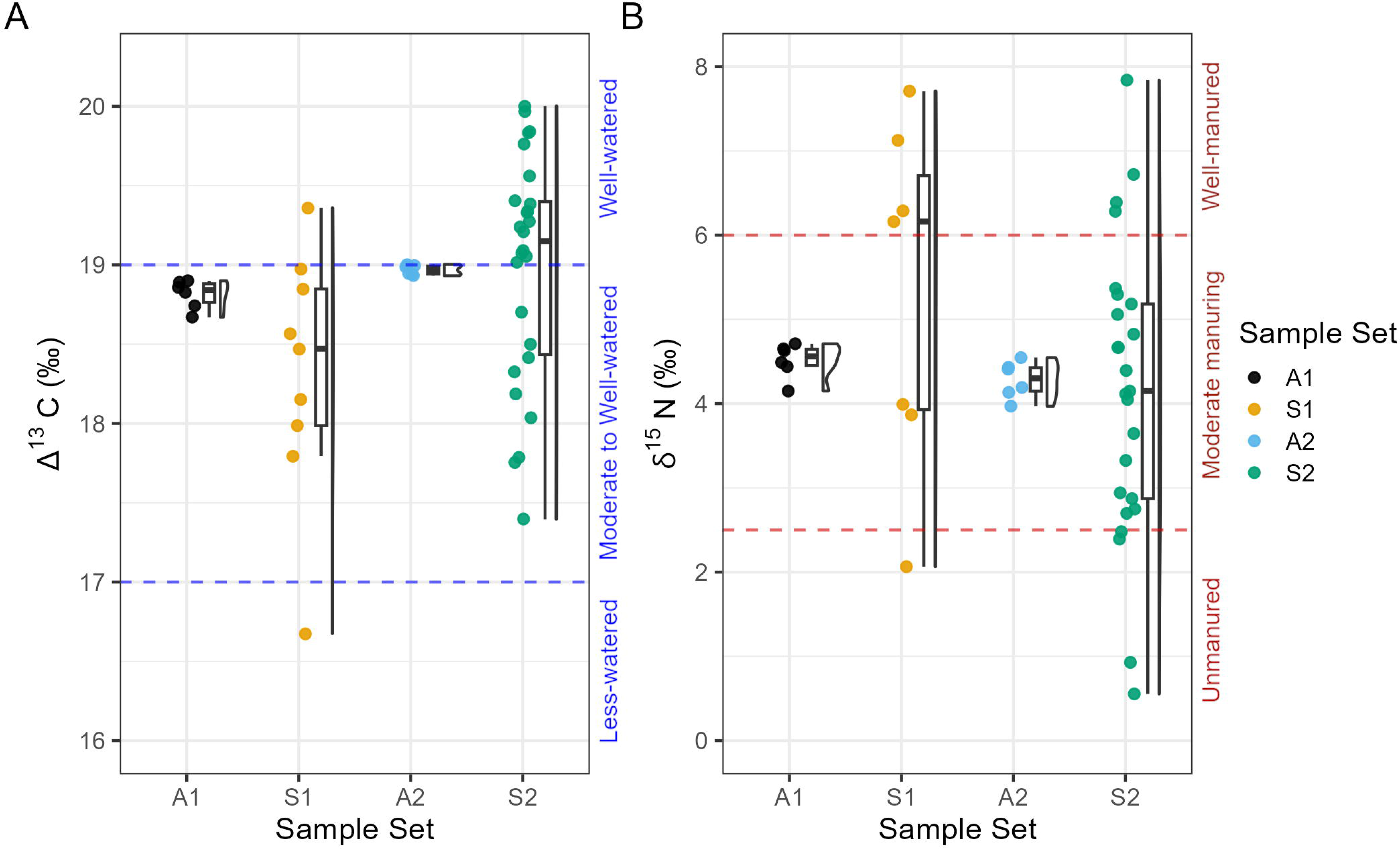
Boxplot of Harappa single-grain and aggregate isotope data. **(A)** Δ^13^C values **(B)** δ^15^N values. Grains with δ^15^N values ≥ 8‰ not shown here (n = 2). **A1:** Aggregate sample of barley (grain n = 10, measurement n = 6). **S1:** Single grains (grain n = 9) **A2:** Aggregate sample consisting of powder from S2 grains (grain n = 26, measurement n = 6) **S2:** Single grain measurements of material aggregated into A2 (grain n = 26).

**Figure 6.**
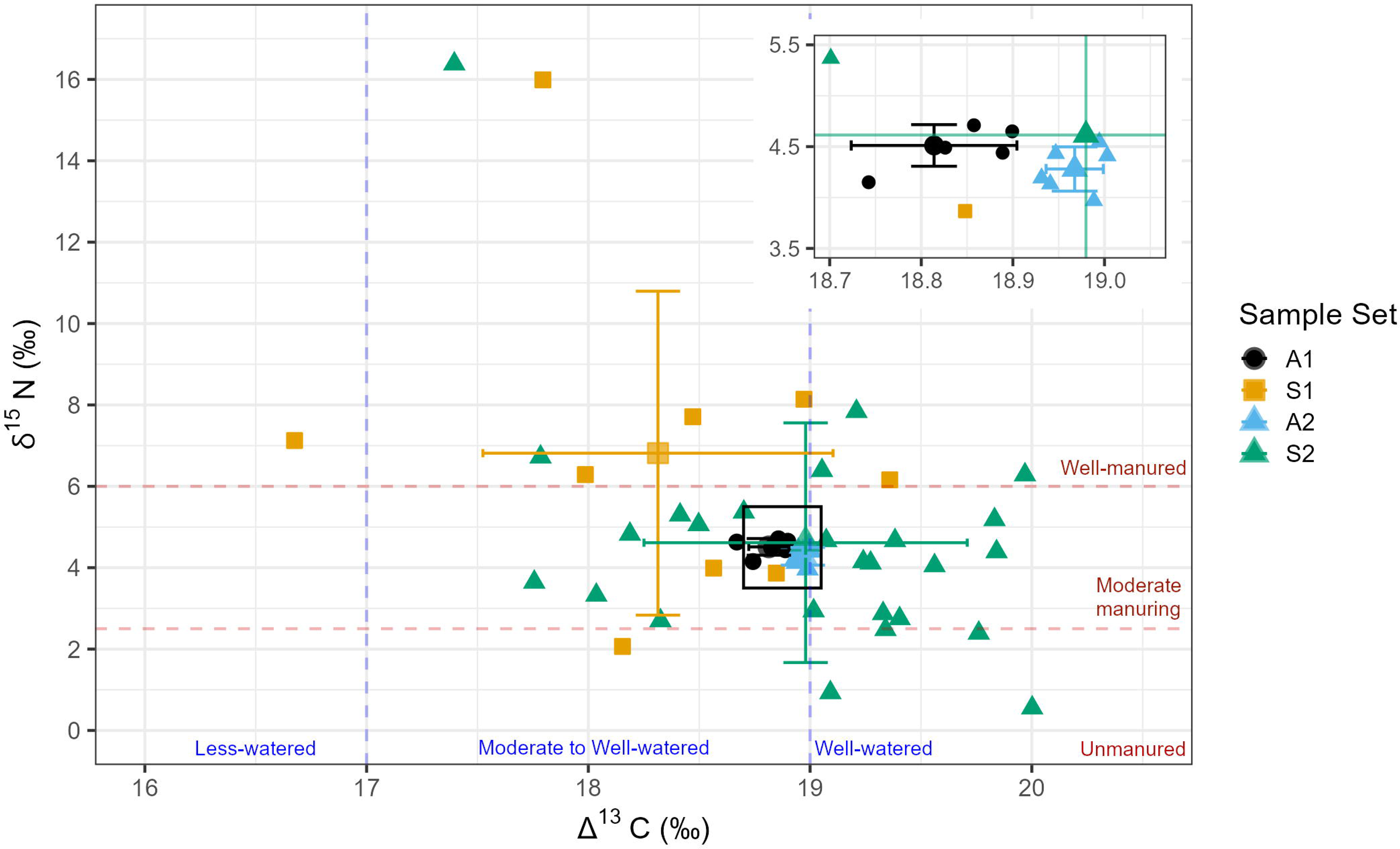
Summary biplot of Harappa single-grain and aggregate Δ^13^C and δ^15^N data. Error boundaries are one standard of deviation from each sample set mean. Inset shows A1 and A2 mean value and standard of deviation. **A1:** Aggregate sample of random grains (grain n = 10, measurement n = 6). **S1:** Single grains (grain = 9). **A2:** Aggregate sample consisting of powder from S2 grains (grain n = 26, measurement n = 6). **S2:** Single grain measurements of material aggregated into A2 (grain n = 26).

**Table 2.** Summary Statistics of Δ^13^C and δ^15^N values for Harappa Trench 42 Barley. **A1:** Aggregate sample of barley (grain n = 10, measurement n=6). **S1:** Single grains (grain n = 9) **A2:** Aggregate sample consisting of powder from S2 grains (grain n = 26, measurement n = 6) **S2:** Single grain measurements of material aggregated into A2 (grain n = 26).

| | *n* | $\delta^{13}\text{C}$<br>*Mean* | $\delta^{13}\text{C}$<br>*SD* | $\delta^{13}\text{C}$<br>*Min* | $\delta^{13}\text{C}$<br>*Max* | $\Delta^{13}\text{C}$<br>*Mean* | $\Delta^{13}\text{C}$<br>*SD* | $\Delta^{13}\text{C}$<br>*Min* | $\Delta^{13}\text{C}$<br>*Max* | $\delta^{15}\text{N}$<br>*Mean* | $\delta^{15}\text{N}$<br>*SD* | $\delta^{15}\text{N}$<br>*Min* | $\delta^{15}\text{N}$<br>*Max* |
| --- | --- | --- | --- | --- | --- | --- | --- | --- | --- | --- | --- | --- | --- |
| **A1** | 6 | -24.8 | 0.1 | -24.8 | -24.6 | 18.8 | 0.1 | 18.7 | 18.9 | 4.5 | 0.2 | 4.2 | 4.7 |
| **S1** | 9 | -24.3 | 0.8 | -25.3 | -22.7 | 18.3 | 0.8 | 16.7 | 19.4 | 6.8 | 4 | 2.1 | 16 |
| **B2** | 6 | -24.9 | 0 | -24.9 | -24.9 | 19 | 0.03 | 18.9 | 19 | 4.3 | 0.2 | 4 | 4.6 |
| **S2** | 26 | -24.9 | 0.7 | -25.9 | -23.4 | 19 | 0.7 | 17.4 | 20 | 4.6 | 2.9 | 0.6 | 16.4 |
| **S1 + S2**<br>*outliers<br>removed | 33 | -24.8 | 0.6 | -25.9 | -22.7 | 18.9 | 0.8 | 16.7 | 20 | 4.5 | 1.9 | 0.6 | 8.1 |
| S1 + S2 | 35 | -24.8 | 0.6 | -25.9 | -22.7 | 18.8 | 0.8 | 16.7 | 20 | 5.2 | 3.3 | 0.6 | 16.4 |

## 5 Discussion

### 5.1 δ^13^C and δ^15^N values in single grains record their local growing conditions in a temperate environment

The Steinzeitpark Dithmarschen (SD) barley exhibited low variation in both carbon and nitrogen isotopes, as expected for C_3_ crops grown in a temperate environment with ample water and no manuring inputs (Figure 2, Figure 3). In C_3_ grasses such as wheat and barley, grain δ^13^C values reflect water availability during crop grain-filling, whether mediated through precipitation, environmental humidity, soil moisture content, or irrigation (Araus et al., 1997a; Cappers and Neef, 2012; Farquhar et al., 1989; Tieszen, 1991; Vogel, 1993). The average δ^13^C value of Steinzeitpark uncarbonized grains (–28.9 ± 0.3‰) corresponds to crops grown in temperate conditions with ample water (“well-watered” = ≤–26‰) (Kohn, 2010; Wallace et al., 2013). There is also low carbon isotope variation (ca. 0.5‰) between inter-panicle mean δ^13^C values. Panicles D and E grains exhibit slightly greater ranges in δ^13^C values than uncarbonized grains from A and B, which is likely due to differences in the sample sizes of uncarbonized grains (Table 1).

The SD single-grain carbon isotope values display extremely low variation across all four panicles. The limited carbon isotope variation displayed in naked barley is consistent with previous studies of single-grain intra-panicle variation in temperate grown wheat, which show ca. ± 0.5‰ variation in grains from a single panicle, and grains between panicles (Heaton et al., 2009). This homogeneity is also consistent with studies showing ca. ± 0.5‰ δ^13^C variation between aggregate samples comprised of grains from crop plants from the same field (Nitsch et al., 2015; Wallace et al., 2015; Jones et al., 2021).

Crop nitrogen stable isotope values are influenced by soil conditions, aridity, and manuring intensity. The frameworks used for interpreting crop δ^15^N values derive from growing experiments where known amounts of manure was applied to fields in arid, semi-arid, and temperate environments δ^15^N values in England, Germany, Morocco, and Syria. Rather than correlating specific amounts of manure to grain nitrogen values, these frameworks rely on estimated benchmarks that gauge the relationship between grain δ^15^N values and manuring practices. Unmanured δ^15^N values fall below ≤ 2.5‰, moderately manured crops range between 2.5 to 6.0‰, and heavily manured crops are approximately ≥ 6.0‰ (Bogaard et al., 2007; Fraser et al., 2011, 2013; Kanstrup et al., 2014; Styring et al., 2016a).

The absence of fertilizer in the SD experimental plot in the form of manure or other amendments means bioavailable nitrogen in the low-nutrient sandy soils are driving plant tissue δ^15^N values (Craine et al., 2015; Szpak, 2014; Denk et al., 2017; Larsson et al., 2024). The low average δ^15^N (0.4 ± 0.6‰) of all uncarbonized grain from the SD plot reflects the lack of direct soil amendments through manuring, and low bioavailable soil N (unmanured δ^15^N = ≤ 2.5‰) (Fraser et al., 2011; Styring et al., 2016a; Larsson et al., 2019). This may be due to increased nitrate leaching in the sandy soils and result in soil nitrogen depleted in ^15^N (Craine et al., 2015; Takebayashi et al., 2010), possibly leading to the low nitrogen isotope values seen here. The uncarbonized grain mean δ^15^N values from Panicles A and B exhibit greater similarity (A, n = 4, 1.2 ±0.1‰; B n = 4, 1.0 ±0.4‰) to each other than with Panicles D (n = 30, 0.7 ± 1.8‰) or E (n = 33, 0.2 ± 0.4‰) from which a larger number of grains were measured. Grain δ^15^N values from Panicle D and E display slightly more variation compared to A and B. In particular, Panicle E displays the lowest mean δ^15^N values, with sixteen grains falling below –0.2‰ (Figure 4) (Amundson et al., 2003; Craine et al., 2009, 2015). The lower nitrogen isotope values of Panicle D and E may be due to slight spatial variation in soil conditions across the cultivation plot, with D and E root systems reflecting particularly low bioavailable nitrogen accessible by each plants root system. The nitrogen isotope values displayed by Panicle D and E barley notably overlap with archaeological barley δ^15^N values from the Funnel Beaker Neolithic site Oldenburg LA77, located ca. 100km away, with δ^15^N values of 0.1‰ (#264 grain n = 10, δ^15^N 0.1‰, #10284 grain n = 15, δ^15^N 0.6‰) (Filipović et al., 2019).

These results underscore the minimal variation in carbon and nitrogen isotope ratios among grains from the same field. Aggregate samples of grain from arid rainfed fields in Jordan, Syria and Spain have demonstrated an ±1‰ difference in aggregate grain δ^13^C values from the same field but harvested in different years, attributed to annual differences in precipitation in arid environments (Flohr et al., 2011, 2019; Wallace et al., 2013). These studies highlight that intra-seasonal variation in precipitation, especially in semi-arid environments, in conjunction with anthropogenic watering practices, significantly influence carbon isotopic variation in charred seeds. In addition, they demonstrate how archaeological grains from contexts that were formed over multiple years, if combined into an aggregate sample, may yield a single carbon isotope ratio that erases key information of annual precipitation or water availability. Nitrogen isotope variation between unmanured grains harvested from single fields in England, Germany, Sweden, and Morocco averages ca. ± 1‰, and increases to ca. ± 2‰ in both manured single grain (Bogaard et al., 2007; Larsson et al., 2019), and manured aggregate samples (Kanstrup et al., 2011; Fraser et al., 2013).

Archaeological studies using single grains have interpreted differences in δ^15^N values of greater than ≥ 2‰ between depositional contexts as either grains from crop plants that experienced different growing conditions in different fields (Lightfoot and Stevens, 2012; Larsson et al., 2019), or disparate access to and application of manure (Larsson et al., 2019). The consistent intra-panicle and inter-panicle isotopic differences found in grains from the same field, as shown here and in other studies demonstrate the interpretive promise of single-grain isotope analysis in archaeological contexts.

### 5.2 Charring has little impact on carbon and nitrogen isotope values

The minimal impact of charring on the carbon and nitrogen isotopic composition of modern seeds from SD confirms previous research demonstrating that charring between 200 to 300°C leaves grains morphology intact and results in a slight ca. 0.1‰ enrichment in ^13^C and ca. 0.3‰ in ^15^N (Fraser et al., 2013; Charles et al., 2015; Nitsch et al., 2015; Stroud et al., 2023a). Grains carbonized at 230°C and 300°C from Panicles A and B exhibit nominally more variation in their nitrogen isotopic compositions than uncarbonized grains (Table 1, Figure). There were slight incremental increases in the mean δ^15^N values of grains carbonized at 230°C and 300°C in both Panicle A (230°C, n = 4, 1.0 ± 0.2‰; 300°C n = 8, 1.5 ± 0.2‰) and Panicle B (230°C, n = 8, 1.2 ± 0.2‰; 300°C, n = 8, 1.5 ± 0.2‰), but no changes in the overall variation (Table 1, Figure 4). In the four grains from Panicle A carbonized at 400°C, only two yielded enough material unrecognizable as grains, for analysis, and these specimens displayed clear deviations from the mean δ^13^C values (n = 2, 26.0 ± 0.8‰) and δ^15^N (n = 2, 4.0 ± 1.2‰). Critically, while there are indications of slightly increased nitrogen isotope variation between grains at higher temperatures, these differences are less than the limited variation displayed between different panicles (Table 1).

### 5.3 Isotope values measured from aggregate sample sets and individual grains reveal different aspects of Harappa agricultural systems

The aggregate and single-grain data from Trench 42 provides two slightly different interpretations of cultivation practices at Harappa ca. 1900 BCE. The high mean aggregate Δ^13^C values (A1 18.8 ± 1.5‰; A2 18.9 ± 1.5‰) for Harappa seeds fall within expectations for well-watered barley (≥ 18.5‰) (Jones et al., 2021; Styring et al., 2016a; Wallace et al., 2013). Single-grain Δ^13^C values show only moderate variation (n = 35, 18.8 ± 0.8‰, range = 16.7 to 20.0‰), with most falling within the approximate range corresponding to medium-to-well-watered barley (Δ^13^C ≥ 17‰), with the exception of a single grain falling into the narrow ‘medium-to-low’ (Δ^13^C 16 to 17‰) watering threshold (Wallace et al., 2013; Flohr et al., 2019; Jones et al., 2021). These values are similar to δ^13^C values of aggregated modern barley from Northwest India grown in flooded fields (n = 101, 18.0 ± 1.5‰), as opposed to rainfed (n = 4, 17.2 ± 2.8‰) or sprinkler-irrigated fields (n = 19, 16.8 ± 1.4‰) (Jones et al., 2021). In addition to the high mean Δ^13^C exhibited by single-grain samples, there is relatively limited carbon isotopic variation, indicating that not only were many of the grains from well-watered fields, but that this water availability was consistently managed (i.e, Styring et al., 2016a) This suggests that at Harappa, water access during the winter growing season was not restricted, as mean carbon discrimination values imply extremely well-watered barley ca. 1900 BCE.

The aggregate mean δ^15^N values (A1: 4.5 ± 0.2‰; A2: 4.3 ± 0.2‰) fall within the benchmarks of low to medium levels of manuring (2.5 to 6.0‰) (Bogaard et al., 2007; Fraser et al., 2011; Styring et al., 2016a), implying that the agricultural soils around Harappa received some manuring amendments, but not in large quantities. In contrast, single-grain δ^15^N values exhibit wide variation (n = 35, 5.2 ± 3.3‰, 0.55 to 16.38‰). Grain nitrogen isotope values are distributed across unmanured (n = 5, ≤2.5‰), medium-to-low (n = 19, 2.5 to 6.0‰), and heavily-manured (n = 11 ≥ 6.0‰) thresholds (Figure 5, Supplemental Information 1) (Fraser et al., 2011, 2013; Larsson et al., 2024; Styring et al., 2016a). While many of the grains span low to moderate manuring conditions, there is also the presence of some grains indicating no manuring, whereas a substantial number of grains indicate highly manured fields. This suggests a wide range of growing conditions and suggests this storage feature held grains from multiple fields subjected to varying manuring practices or access to manure.

### 5.4 Aggregate or single-grain stable isotope analysis impacts interpretation of plant management practices

The relationships between crop stable carbon and nitrogen values and agricultural organization hinges on the correlation of manuring and watering inputs to varying labor inputs and land use (Bogaard, 2004; Styring et al., 2017a; Bogaard et al., 2019). The interpretative framework for archaeobotanical stable isotope data guiding this correlation relies on models of labor- and land-limited agriculture derived from ethnographic studies of traditional smallholder farmers in Africa and Europe (Netting, 1993; Halstead, 2014). Labor-limited intensive practices are focused on small households dependent solely on scarce household labor for field management (Bogaard, 2004, 2005; Halstead, 2014). The expectation is that an increase in production under labor-limited conditions would entail high watering and high manuring inputs concentrated into small plots by nuclear households (Bogaard, 2005). These agricultural practices would be reflected in the isotopic composition of cultivated plants visible as high crop Δ^13^C and δ^15^N values (Wallace et al., 2013; Styring et al., 2016a). Land-limited extensive practices are associated with the use of plow agriculture, increasing available labor through traction to bring more land under cultivation, but the attendant spatial extensive land use means there are fewer watering, manuring, and labor inputs per field (Styring et al., 2017a). These less intensive applications of water and manuring would translate to a decrease in Δ^13^C and δ^15^N values (Styring et al., 2016a).

This framework traces connections between stable nitrogen and carbon isotope values that may indicate changing agricultural production that shift from intensive to extensive practices (Bogaard et al., 2013, 2019; Styring et al., 2017b, 2017a; Yang et al., 2022). To date, exploration of land vs. labor limited agriculture in ancient cultivation systems has relied on isotope values derived from aggregate grain samples, stating their use for large-scale regional and temporal comparisons. Aggregate samples containing large numbers of seeds attempt to ensure that samples encompass the variation of growing conditions in a field (Nitsch et al., 2015; Bogaard et al., 2016). Though archaeological specimens could span multiple growing seasons and thus different growing conditions (Riehl, 2020), aggregate studies use this possibility of wide ranging conditions to obtain isotopic information of the range of possible cultivation practices at a site, making large scale comparisons viable (Kanstrup et al., 2012; Styring et al., 2017a).

The Trench 42 aggregate samples (A1 and A2) mean Δ^13^C values correspond to those expected for moderately to well-watered crops (Jones et al., 2021), while the mean δ^15^N values fall within the benchmarks of medium to potentially low levels of manuring (Bogaard et al., 2007; Fraser et al., 2011; Styring et al., 2016a). Within the frameworks of labor- and land-limited agriculture, in conjunction with the widespread evidence of traction and plow agriculture at Harappa, this data suggests limitations on available land as opposed to available labor. It may be that labor was abundant and the cultivation of additional fields gave rise to land becoming more scarce and increasingly valuable (Bogaard, 2005). Therefore, within this framework, the isotope values from aggregate sample broadly correspond to evidence for land-limited extensive practices at Harappa (Bogaard, 2004; Bogaard et al., 2019).

The overlap between the mean Δ^13^C and δ^15^N values of single-grain S1 and S2 and aggregates A1 and A2 indicate that an aggregated sample pool captures representative signature of cultivation. However, the variation displayed in S2 compared to A2 illustrates how isotope values measured from aggregate samples do not fully describe the full range of isotopic variation expressed in archaeological grains that better identifies diversity in ancient agricultural practices. Along these lines, the carbon and nitrogen isotopic variation shown by individual barley grains (S1 and S2) from Trench 42 complicates an interpretation of land-limited production (Figure 5, 6). The mean Δ^13^C and δ^15^N values produced by S1 and S2 are similar to A1 and A2, indicating that the aggregate samples capture an average isotopic variation in a primary context (Figure 5). However, the variation in single-grain δ^15^N values from Trench 42 barley (δ^15^N 5.2 ± 3.3‰) imply the presence of both intensively manured fields and fields receiving little to no manure (Figure 6). On the other hand, the variation in single-grain Δ^13^C values indicates consistent watering practices or conditions with well-watered barley (Δ^13^C 18.8 ± 0.8‰). Wide ranges of carbon and nitrogen isotope variation resulting from different field conditions might be expected from extensive agricultural practices that bring more land under cultivation (Styring et al., 2017a). However, this pattern of limited Δ^13^C and wide δ^15^N variation at Harappa suggests both active water management of barley, and concurrent practices that may represent both low and highly manured fields. These results underscore the need to further evaluate the variation present in both intensive and extensive agricultural systems. Greater inter-grain isotopic variation could be reasonability expected when more extensive agricultural practices are used as more land, which would in likely encompass a greater variety of growing conditions at local scales, might be plausibly engaged by ancient agriculturalists. Additional exploration of the relationship between spatially dispersed cultivation practices and isotopic variation expressed in grain assemblages is required.

## 6 Conclusion: Promise and limitations for single grain isotope analyses

If a depositional context contains grains from multiple fields, a wide geographic area, or a wide temporal scale, an aggregate sample of ten grains will encompass more isotopic variation than expected for a single set of growing conditions as defined by analysis of modern aggregate samples (Nitsch et al., 2015; Stroud et al., 2023a). Recent recommendations suggest aggregate samples are more appropriate for primary depositional contexts that may be presumed to represent single years or growing conditions, as this is less likely to collapse isotopic variation representing multiple fields or harvests. When sampling a secondary depositional context, such as a bulk stratigraphic soil sample or tertiary redeposited midden, to compensate for the potential loss of isotopic information representing multiple fields or harvests, 2–3 aggregate samples of ∼10 grains each should be selected for stable isotope analysis (Vaiglova et al., 2023). This analysis from a Harappa depositional storage context suggests that the average created by aggregate samples is unlikely to encompass those ranges of variation.

The sampling of individual grains for stable isotope analysis is subject to several clear restraints. Limited funding resources, time, or labor may restrict researchers’ abilities to measure larger number of individual grains that would fully document the complete range of isotopic variation represented in an assemblage. Sample screening protocols also impact sampling strategy. Establishing if grain specimens contain humic or fulvic acids, carbonates, or other contaminants from the burial environment, and subsequent determination of wet chemistry pre-treatment protocols, requires for Attenuated Total Reflectance Fourier Transform Infrared Spectrometry (ATR-FTIR) approximately 1–2mg of charred sample (Vaiglova et al., 2014; Brinkkemper et al., 2018). Although the amount of sample demands for ATR-FTIR instruments are low, requiring a very small volume of powder to cover the ATR window, single small grains may be consumed by this process. Larger specimens, even if sub-sampled for pre-screening with sufficient material for wet chemistry, may experience more extensive sample loss during the pretreatment phase. For example, mass loss of aggregated powers (n_grain_ = 5) exposed to 0.5 HCl at 80 °C for 30 minutes followed by three rinses with Milli-U water to remove nitrate, humic, or carbonate contaminates, results in ca. 60% sample loss (Vaiglova et al., 2014).

Environmental context can also play into decisions whether or not to employ an aggregate or single seed sample strategy. The measurement of individual grains for δ^13^C is especially important in water-limited environments where variation in watering practices and irrigation systems, crop emergence and ripening time, and landscape dependent soil moisture levels in agricultural plots would impart marked differences in the carbon isotopic composition of cereal crops (Wallace et al., 2013; Styring et al., 2016a; Flohr et al., 2019). Single-grain sampling may be less important in well-watered, temperate environments where carbon isotope variation at the floral base of the food web is less pronounced (Heaton, 1999; Kohn, 2010; Diao et al., 2023).

Despite the promise of the high-resolution isotopic information gained through single-seed analyses, it is important to keep in mind that carbon and nitrogen isotope values measured from aggregated samples describe mean isotope values averaged from single grains, thus providing a broad picture of plant husbandry strategies used at ancient settlements. Whether or not it is mandatory to analyze single seeds for carbon and nitrogen isotopes depends on the research question at hand (Gron et al., 2021; Gavériaux et al., 2022; Vaiglova et al., 2023). To further test the viability of single-grain stable isotope analysis, future studies should assess ranges of variation present in archaeobotanical stable isotope data between different sites, context types, and through time. Studies that rely on large, regional-scale comparative datasets to address diachronic change in extensification/intensification in relation to landscape urbanization, emergent staple finance systems, or climatic change might be best served by an aggregate sampling strategy that would establish broad, settlement level agricultural systems. Tracing intra-settlement diversity in cultivation practices at the household or neighborhood level, or at urban sites such as Harappa, might be better achieved through a single-grain sampling strategy that closely queries intra-contextual variability.

The overlap in single-grain and aggregate sample means demonstrate the effectiveness of previous studies investigations into adequate sample sizes for archaeobotanical isotope analysis (Stroud et al., 2023b; Nitsch et al., 2015). This overlap between sample sets, in addition to the fully represented variation in single-grain isotope values, offers a path to explore the isotopic variation present in archaeobotanical assemblages, and elucidate our understanding of the cumulative outcomes of farmers decision making amidst dynamic environmental cultural contexts.

## Acknowledgements

This research was funded by the Deutsche Forschungsgemeinschaft (DFG, German Research Foundation) EXC ROOTS Excellence Cluster (DFG EXC 2150). We thank Yasmin Dannath for facilitating access to the archived modern grain samples from Steinzeitpark Dithmarschen. We thank Fiona Friedrich-Walker for laboratory assistance and brainstorming, Damini Pant for initial comments, and Steve Weber for his insights into the Harappa archaeobotanical collection.

## Credit

NFJ: Conceptualization, Formal analysis, Investigation, Validation, Visualization, Writing – original draft, Writing – review & editing. CWS: Writing – Formal analysis, Data Curation, review and editing, MK: Resources, Writing – review & editing. JDG: Supervision, Writing – review & editing. CMAK: Conceptualization, Funding acquisition, Resource, Supervision, Writing – original draft, Writing – review & editing.

